# End-to-end deep image reconstruction from human brain activity

**DOI:** 10.1101/272518

**Authors:** Guohua Shen, Kshitij Dwivedi, Kei Majima, Tomoyasu Horikawa, Yukiyasu Kamitani

## Abstract

Deep neural networks (DNNs) have recently been applied successfully to brain decoding and image reconstruction from functional magnetic resonance imaging (fMRI) activity. However, direct training of a DNN with fMRI data is often avoided because the size of available data is thought to be insufficient to train a complex network with numerous parameters. Instead, a pre-trained DNN has served as a proxy for hierarchical visual representations, and fMRI data were used to decode individual DNN features of a stimulus image using a simple linear model, which were then passed to a reconstruction module. Here, we present our attempt to directly train a DNN model with fMRI data and the corresponding stimulus images to build an end-to-end reconstruction model. We trained a generative adversarial network with an additional loss term defined in a high-level feature space (feature loss) using up to 6,000 training data points (natural images and the fMRI responses). The trained deep generator network was tested on an independent dataset, directly producing a reconstructed image given an fMRI pattern as the input. The reconstructions obtained from the proposed method showed resemblance with both natural and artificial test stimuli. The accuracy increased as a function of the training data size, though not outperforming the decoded feature-based method with the available data size. Ablation analyses indicated that the feature loss played a critical role to achieve accurate reconstruction. Our results suggest a potential for the end-to-end framework to learn a direct mapping between brain activity and perception given even larger datasets.

## 1. Introduction

Decoding visual contents from brain activity using deep neural networks (DNN) has recently shown promising results. DNNs have been applied on functional magnetic resonance imaging (fMRI) data to reconstruct the perceived image (Güçlütürk et al., 2017; Shen et al., 2017; Han et al., 2017; Seeliger et al., 2017) or to identify the object category (Horikawa and Kamitani, 2017). However, the above methods avoid directly training a DNN model on the fMRI data due to smaller dataset sizes in fMRI studies. To solve the dataset size issue, the feature representation from a DNN pretrained on a large scale image dataset was used as a proxy for the neural representations of the human visual system. Hence, these methods involve two independent steps, 1) decoding DNN features from fMRI activity and 2) reconstruction or identification using the decoded DNN features.

In image generation studies in computer vision, a DNN can be trained using an end-to-end approach to directly generate the image from a modality correlated with the images, e.g., captions (Mansimov et al., 2015) and DNN features (Dosovitskiy and Brox 2016a, 2016b). In the end-to-end approach, the DNN model generates the image directly from the correlated modality. However, the dataset sizes used to train such models are usually larger as compared to the dataset sizes in fMRI studies. For instance, Mansimov et al. (2015) trained a caption-to-image model on Microsoft COCO dataset that consists of 82,783 images, each annotated with at least 5 captions. Dosovitskiy and Brox (2016a) trained a DNN model on ImageNet training dataset (over 1.2 million images) to reconstruct images from corresponding DNN features. On the other hand, the largest fMRI dataset used for reconstruction in Shen et al. (2017) consists of only 6,000 training samples. Thus, training a DNN to reconstruct images directly from fMRI data is often avoided and considered infeasible due to the smaller dataset size.

In this study, we sought to evaluate the potential of the end-to-end approach to obtain a direct mapping from fMRI activity to stimulus space given a limited training dataset. Training a DNN in an end-to-end manner implies that the input to the DNN is the fMRI activity and output of the DNN is the reconstruction of the perceived stimulus. If we can successfully perform reconstruction using the end-to-end approach, then we can avoid feature decoding step used in earlier studies and reconstruct directly from the fMRI activity.

For designing an end-to-end DNN model to reconstruct from fMRI data, we transformed the fMRI data into a 1-dimensional vector and therefore, the reconstruction model must transform 1-dimensional fMRI data to a 3-dimension image in RGB color space. The neural network architectures that transform 1-dimensional image features to the original image are thus well-suited for this purpose. The fMRI data from the visual cortex can also be considered as a representation of the perceived image, and thus can be used as an input in such architectures for the end-to-end training.

Dosovitskiy and Brox (2016a) proposed a DNN based method to generate the original image from the corresponding features by optimizing in image space. The loss in image space only is usually insufficient to obtain a good reconstruction since it generates a reconstruction that is an average of all the possible reconstructions with the same distance in image space and hence the reconstructed images are blurred. The feature loss in high dimensional space, also called perceptual loss, constrains the reconstruction to be perceptually similar to the original image. Adversarial loss (Goodfellow et al., 2014) constrains the distribution of the reconstructed images to be close to the distribution of natural images. In a subsequent study, Dosovitskiy and Brox (2016b), showed that reconstruction from features could be improved by introducing feature and adversarial loss terms. Hence, we adopted the approach of Dosovitskiy and Brox (2016b) to reconstruct the perceived stimuli directly from the fMRI activity. We modified their model to take input directly from the fMRI activity and trained the model from scratch on the dataset from Shen et al. (2017).

In this study, we first demonstrate that we can obtain reconstructions resembling the original stimulus images from the model trained on this dataset. We further explore the generalizability of the proposed method on artificial shapes and alphabetical letters. To understand the effect of training dataset size on reconstruction quality, we compare the reconstruction results as the training dataset size gradually increased from 120 samples to 6,000 samples. Finally, to investigate the effects of the different loss functions used in the reconstruction method, we perform an ablation study by removing one loss function at a time and performing a subjective and objective comparison of the reconstruction results.

## 2. Materials and Methods

In this section, we briefly describe the method we used for our experiments and the details of the dataset. For more details regarding the image reconstruction method please refer to Dosovitskiy and Brox (2016b), and for details regarding dataset, please refer to Shen et al. (2017).

### 2.1. Problem Statement

Let **X** ∈ ℝ^*w*×*h*×3^ be the stimulus image displayed in the perception experiment where *w* and *h* are width and height of the stimulus image respectively and 3 denotes the number of channels (RGB) of the color image. Let **f** ∈ ℝ^*N*^ be the corresponding preprocessed fMRI vector for the brain activity recorded during the perception and *N* is the number of voxels in the visual cortex (VC). We are interested in obtaining a reconstruction of the stimulus from fMRI vector **f**.

To solve this problem, we use a DNN **G**_θ_ with parameters **θ** which performs non-linear operations on **f** to obtain a plausible reconstruction **G_θ_**(**f**) of the stimulus image.

### 2.2. Image reconstruction model

We modified the DNN model proposed by Dosovitskiy and Brox (2016b) to reconstruct stimulus images from fMRI data.

For the fMRI vector **f** corresponding to the stimulus image **X**, the model is trained to generate a plausible reconstruction image **G_θ_**(**f**) from **f**. The network architecture (Figure 1A) consists of three convolutional neural networks: a generator **G_θ_** which transforms the fMRI vector **f** to **G_θ_**(**f**), a discriminator **D_Φ_** which discriminates the reconstructed image **G_θ_**(**f**) from the natural image **X**, and a comparator **C** which performs the comparison between the reconstructed image **G_θ_**(**f**) and the original stimulus image **X** in the feature space.

**Figure 1.**
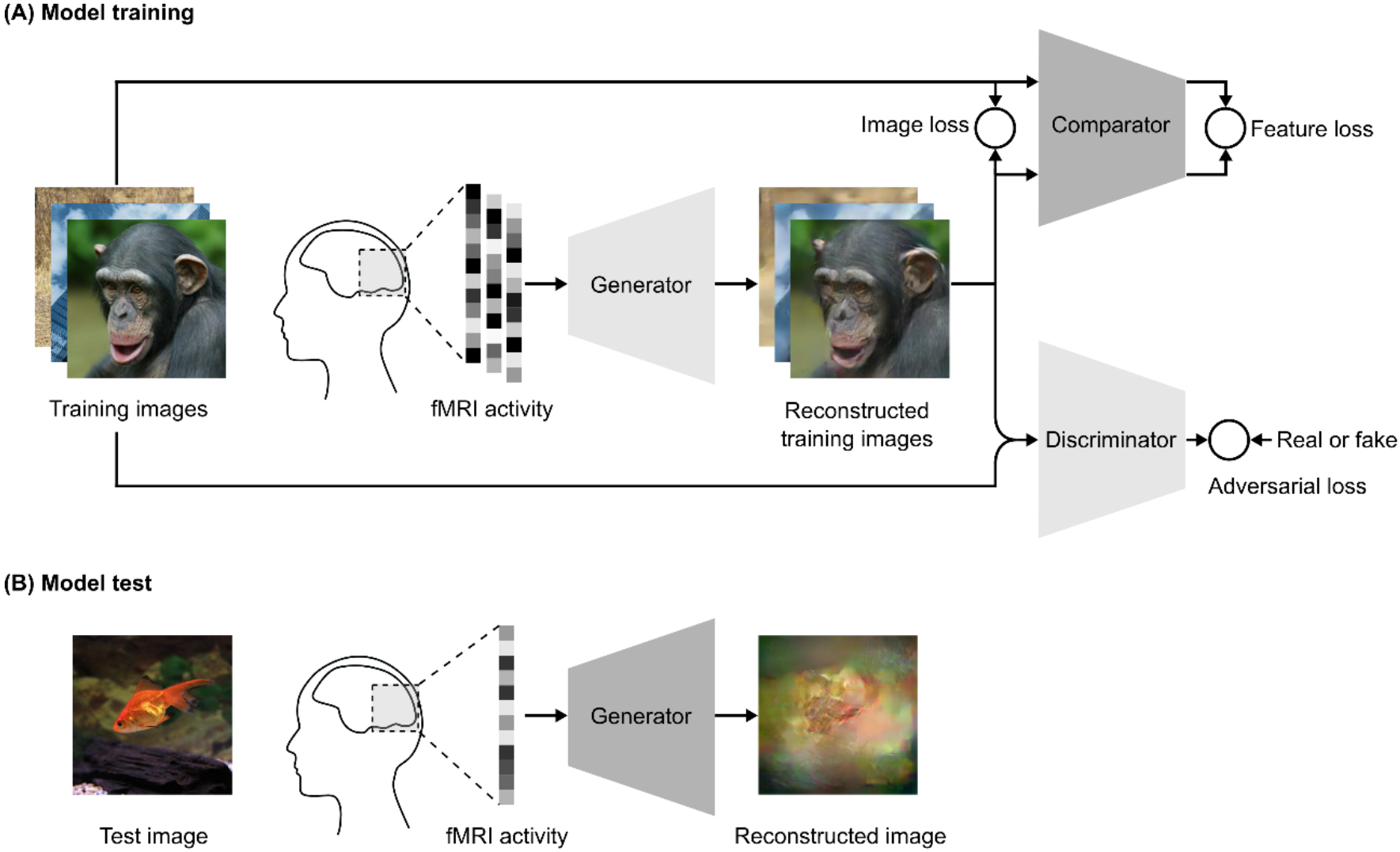
Schematics of our reconstruction approach. **(A) Model training.** We use an adversarial training strategy adopted from Dosovitskiy and Brox (2016b), which consists of 3 DNNs: a generator, a comparator, and a discriminator. The training images are presented to a human subject, while brain activity is measured by fMRI. The fMRI activity is used as an input to the generator. The generator is trained to reconstruct the images from the fMRI activity to be as similar to the presented training images in both pixel and feature space. The adversarial loss constrains the generator to generate reconstructed images that fool the discriminator to classify them as the true training images. The discriminator is trained to distinguish between the reconstructed image and the true training image. The comparator is a pre-trained DNN, which was trained to recognize the object in natural images. Both the reconstructed and true training images are used as an input to the comparator, which compares the image similarity in feature space. **(B) Model test.** In the test phase, the images are reconstructed by providing the fMRI activity of the test image as the input to the generator.

The input to the generator is the fMRI vector **f** from VC and the output is the reconstructed image **G_θ_**(**f**). The generator consists of three fully connected layers followed by six upconvolution layers to generate the final reconstruction image **G_θ_**(**f**).

The comparator network **C** is Caffenet trained on Imagenet dataset for the image classification task. The Caffenet model is a replication of Alexnet (Krizhevsky et. al. 2012) model with the order of pooling and normalization layers switched and without relighting data-augmentation during training. The network consists of 5 convolutional and 3 fully connected layers. We used the last convolutional layer of the comparator to compare the reconstructed image and the original stimulus image in feature space. The parameters of the comparator were not updated during the training of the reconstruction model.

The discriminator **D_Φ_** consists of five convolutional layers followed by an average pooling layer and two fully-connected layers. The output layer of the discriminator is a 2-way softmax and the network is trained to discriminate the original stimulus image from the reconstructed image. The purpose of the discriminator is to learn to differentiate between original stimulus images and images reconstructed by the generator. The generator is trained concurrently to optimize an adversarial loss function to fool the discriminator into classifying the reconstructed image as the real stimulus image. The adversarial loss forces the generator to generate more realistic images closer to image distribution of the training data.

The architectures of the discriminator and comparator networks were the same as in Dosovitskiy and Brox (2016b). The generator was modified to take its input from the fMRI data as opposed to DNN features in Dosovitskiy and Brox (2016b).

Let **X**_*i*_ denote the *i^th^* stimulus image in the dataset, **f**_*i*_ denote the corresponding fMRI data for the *i^th^* image and **G_θ_**(**f**_*i*_) denote the reconstruction output of the generator. The parameters **θ** of generator **G_θ_** are updated to minimize the weighted sum of three loss terms for a minibatch using stochastic gradient descent: loss in image space *L*_img_, feature loss *L*_feat_, adversarial loss *L*_adv_:

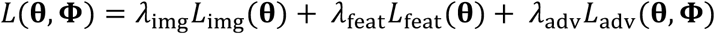

where

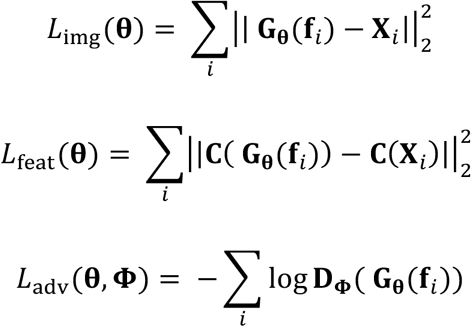

and *λ*_img_, *λ*_feat_, and *λ*_adv_ denote the weights of the loss in image space *L*_img_, feature loss *L*_feat_, and adversarial loss *L*_adv_, respectively.

The discriminator is trained concurrently with the generator to minimize *L*_discr_

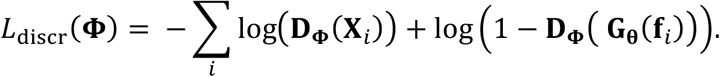

The parameters of the comparator **C** are fixed throughout the training since it is used just for the comparison in feature space and therefore no update was required.

We performed the training using Caffe framework (Jia et al., 2014) and modified the implementation of the model provided by Dosovitskiy and Brox (2016b). The weights of generator and discriminator were initialized using MSRA (He et al., 2015) initialization. The comparator weights were initialized by Caffenet weights trained on Imagenet classification. We used Adam solver (Kingma and Ba, 2015) with momentum *β*_1_ = 0.9, *β*_2_ = 0.999 and initial learning rate 0.0002 for optimization. We used a batch size of 64 and trained for 500,000 mini-batch iterations in all the experiments. Following the similar training procedure as Dosovitskiy and Brox (2016b), we temporarily stopped updating the discriminator if the ratio of *L*_discr_ and *L*_adv_ was below 0.1. This was done to prevent the discriminator from overfitting. The weights of the individual loss functions *λ*_img_, *λ*_feat_, and *λ*_adv_ were set to *λ*_img_ = 2*e* − 6, *λ*_feat_ = 0.01, and *λ*_adv_ = 100.

We applied image jittering during the training for data augmentation and to take into account the eye movement of the subjects during image presentation experiment. Generally, for a typical subject, the size of eye movement is about 1 degree viewing angle. The viewing angle for the presented images is 12 degrees. All the training images are resized to 248 × 248 before the training. During the training, we randomly cropped a 227 × 227 window from each training image as the target image for each iteration to ensure that the largest jittering size was 12 * (248 − 227)/227~1 degrees.

For dataset size analysis we trained the reconstruction model with variable number of training samples for 100 epochs with a batch size of 60. The rest of the hyperparameters were same as the previous analysis.

### 2.3. Dataset from (Shen et al., 2017)

We used an fMRI dataset from a previous reconstruction study (Shen et al., 2017). This dataset was used to reconstruct stimulus images from visual features of a deep convolutional neural network decoded from the brain.

The stimulus images in the dataset were categorized into four types: training-natural images, test-natural images, artificial shapes and alphabetical letters. The natural images used for the experiment were selected from 200 representative object categories in the ImageNet (Deng et al., 2009; 2011, fall release) dataset. The training-natural image dataset consisted of 1,200 images from 150 object categories and test-natural image dataset consisted of 50 images from 50 object categories. There was no overlap between the categories used in the training and test datasets. The artificial shapes consisted of 40 images obtained by combining 8 colors and 5 shapes. The alphabetical letters consisted of 10 images of different letters from the English alphabet in black color.

The image presentation experiments consisted of four distinct types of sessions corresponding to the four categories of stimulus images described above. In one set of the training-natural image session, a total of 1,200 images was presented only once. This set of training session was repeated five times. In the test-natural image session, the artificial-shape session, and the alphabetical letter session, 50, 40, and 10 images were presented 20, 20, and 12 times each, respectively. The presentation order of the images was randomized across runs.

The fMRI data obtained during the image presentation experiment were preprocessed for motion correction followed by co-registration to the within-session high-resolution anatomical images of the same slices and subsequently to T-1 anatomical images. The coregistered data were then re-interpolated as 2 × 2 × 2 mm voxels.

The fMRI data samples were created by first regressing out nuisance parameters, including a linear trend, and temporal components proportional to six motion parameters calculated by the SPM5 (http://www.fil.ion.ucl.ac.uk/spm) motion correction procedure, from each voxel amplitude for each run. After that, voxel amplitudes were normalized relative to the mean amplitude of the initial 24-s rest period of each run, and were despiked to reduce extreme values (beyond ±3SD for each run). The voxel amplitudes were then averaged within each 8-s (training natural image-sessions), 12-s (test natural-image, artificial-shapes, and alphabetical-letter sessions) stimulus block (four or six volumes), after shifting the data by 4 s (two volumes) to compensate for hemodynamic delays.

The voxels used for the reconstruction were selected from the visual cortex (VC), which consisted of lower visual areas: V1, V2, V3, and V4 and higher visual areas: lateral occipital complex (LOC), fusiform face area (FFA), and parahippocampal place area (PPA). V1, V2, V3, and V4 were identified using a retinotopy experiments and LOC, FFA, and PPA were identified using a functional localizer experiments (Shen et al., 2017).

The fMRI data was further normalized before using it as an input to the reconstruction model. The mean 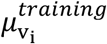 and standard deviation 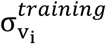 of amplitude a_v_i__ of each voxel v_i_ in VC were estimated across the training data. Then for both training and test data, the normalized amplitudes 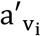 of voxel v_i_ ∈ *VC* for each sample were estimated as follows:

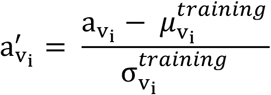

To compensate the statistical difference between the training and test fMRI data (we performed trial-averaging for the test fMRI data while we considered each trial as an individual sample for the training fMRI data), we rescaled the test fMRI data by a factor of 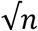 where *n* is number of trials averaged, before we use the test fMRI data as the input to the generator.

For dataset size analysis, initially a fixed number of training images and the corresponding fMRI activity from five trials were selected for training. As the dataset size was increased more training images with fMRI activity were subsequently added. The training dataset size was increased gradually from (5 × 24)120 to (5 × 1200) 6,000 training samples.

### 2.4. Evaluation

We evaluated the quality of reconstruction using both objective and subjective assessment methods. For both assessment methods, we performed a pairwise similarity comparison, where one reconstructed image was compared with two possible candidate images, one was the original stimulus image from which the reconstruction was obtained and the other was randomly selected from the test dataset of the same image type.

For the subjective assessment, we conducted a behavioral experiment similar to Shen et al. (2017). In the experiment, another group of 13 raters (6 females and 7 males, aged between 19 and 48 years) were presented a reconstructed image with two possible candidate images and were asked to select the candidate image which appears more similar to the reconstructed image.

For the objective assessment, we compared the pixel-wise correlation coefficients of the reconstructed image with the two candidate images and selected the candidate image with the higher correlation coefficient.

For both assessments, we calculated the percentage of trials in which the original stimulus image was selected and used it as the quality measure. The trial for each reconstructed image was conducted with all the pairs of the images from same type.

We conducted another behavioral experiment to study the effect of different loss terms in the proposed approach. 3 raters (1 female and 2 males, aged between 30 and 37 years) from a different group were presented one original stimulus image and two reconstructed images generated from different combination of loss terms. The raters were asked to judge which one of the reconstructions showed higher resemblance with the original stimulus image. This pairwise comparison was conducted for 8 pairs of combinations of loss terms for all the stimulus image in test dataset. We use the winning percentage as the quantitative measure to compare reconstructions obtained from different combinations of loss terms. The winning percentage is the percentage of trials in which the reconstruction from one combination was judged better than the other. For computing the winning percentage from pixel-wise correlation coefficients, the reconstruction with higher correlation coefficient was selected. For more details about the design of both the behavioral experiments, please refer to Shen et al. (2017).

## 3. Results

### 3.1. Image reconstruction

We trained the reconstruction model on the training session samples of the fMRI dataset from Shen et al. (2017) which consisted of perception fMRI data corresponding to 1,200 natural images. In the training session, each stimulus image was presented to the subject 5 times. We treated each stimulus presentation as a separate training sample for the reconstruction model. Therefore, the training dataset we used, consisted of 6,000 (5 × 1200) samples.

We evaluated the reconstruction quality on three test datasets: natural images, artificial shapes and alphabetical letters. For generating reconstructions, fMRI samples corresponding to the same image (20 samples in the test-natural image session, 20 samples in the artificial shapes session, and 12 samples in the alphabetical letters session) were averaged across trials to increase the signal to noise ratio, averaged fMRI samples were used as input to the trained generator. Figure 2A shows some example images from the test natural images and their corresponding reconstructions from three different subjects obtained using our model. The reconstructions from all three subjects closely resemble the natural -image stimuli in shape. The color, however, is not preserved in some of the reconstructions. The reconstruction results from our model show that despite utilizing a small dataset, it was possible to train a model from scratch that could reconstruct visually similar images from fMRI data with high accuracy (Figure 2B; 78.1% accuracy by pixel-wise spatial correlation, 95.7% by human judgment).

**Figure 2.**
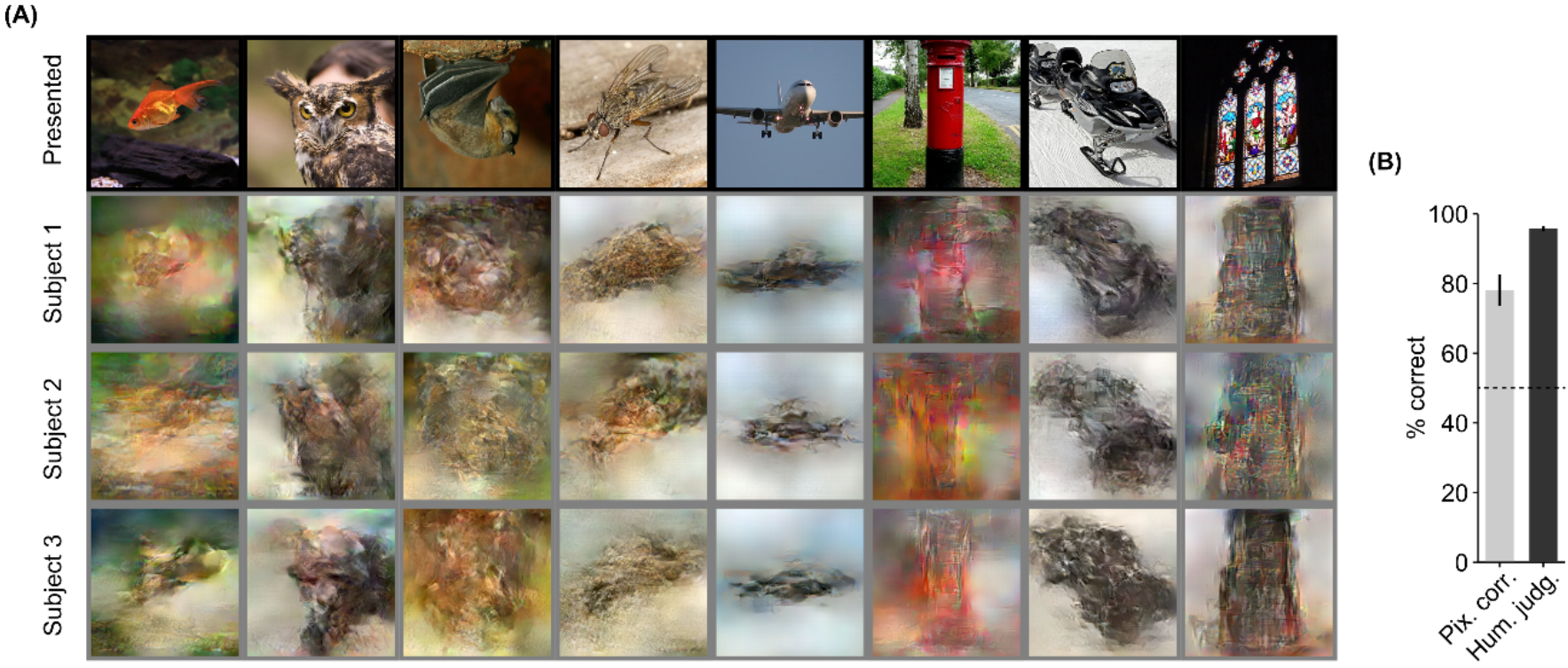
Results of reconstruction of natural images. **(A)** Presented and reconstructed natural images are shown here. The presented images (in black frames) are shown in the top row. Three corresponding reconstructed images (in gray frames) from each of the three subjects are shown underneath. **(B)** Reconstruction accuracy of natural images in terms of percentage of correct pair-wise classification based on both pixel correlation and human judgment (error bars, 95% confidence interval (CI) across samples; three subjects pooled; chance level, 50%).

Further, we evaluated the generalization of the method using artificial shapes similar to Shen et al. (2017). We demonstrate that using the proposed approach, artificial shapes (Figure 3A) can be reconstructed with high accuracy (Figure 3B. 69.3% by pixel-wise spatial correlation, 92.7% by human judgment) even though the model was trained on natural images.

**Figure 3.**
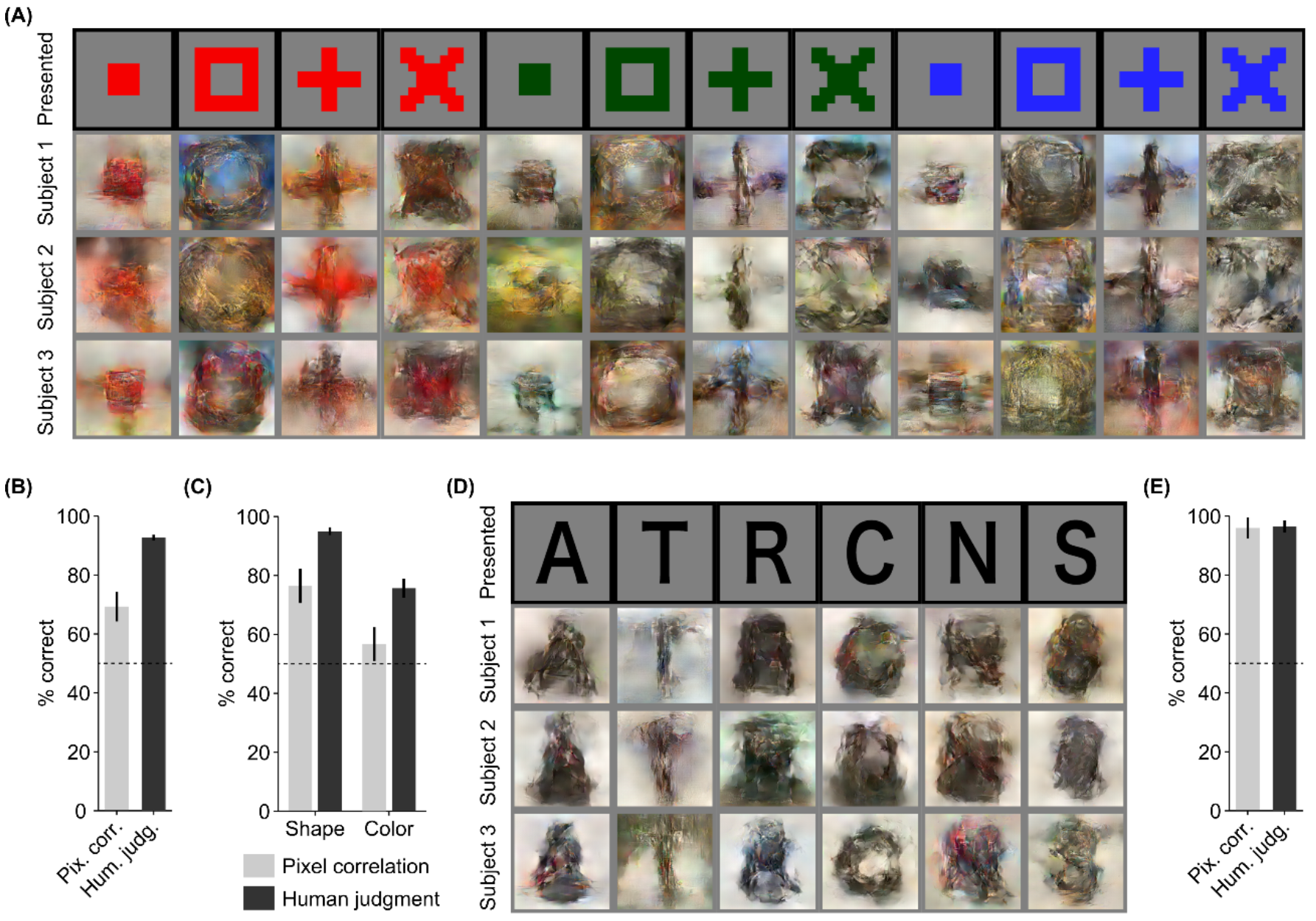
Reconstruction of artificial shapes and alphabetical letters. **(A)** Reconstruction of artificial shapes. The original stimulus images (in black frames) are shown in the top row. Three corresponding reconstructed images (in gray frames) from each of the three subjects are shown underneath. **(B)** Reconstruction accuracy of artificial shapes. **(C)** Reconstruction accuracy of both shape and color. **(D)** Reconstruction of alphabetical letters. **(E)** Reconstruction accuracy of alphabetical letters. For **(B)**, **(C)** and **(E)**, reconstruction accuracy is assessed in terms of percentage of correct pair-wise classification based on both pixel-wise correlation and human judgment (error bars, 95% CI across samples; three subjects pooled; chance level, 50%).

From the artificial shape reconstruction results, we observed that the shape of the stimulus is well preserved in the reconstructions. However, the color in the reconstructions is preserved only for the red-colored shapes, while the reconstructions of the other-colored shapes do not show resemblance in terms of color. To compare the reconstruction quality in terms of shape and color, we performed comparison across the reconstructed images of same shapes and colors. The quantitative results from Figure 3C (shape: 76.5% by pixel-wise spatial correlation, 95.0% by human judgment, color: 56.7% by pixel-wise spatial correlation, 75.6% by human judgment) suggest that reconstructed images show more resemblance in terms of shape as compared to color.

We further show the generalizability of our approach by showing highly accurate reconstructions of the alphabetical letters images (Figure 3D). The alphabetical letters reconstruction accuracy was 95.9% according to pixel-wise spatial correlation, and 96.4% according to human judgment.

We compared the reconstruction accuracy of the proposed method with Shen et al. (2017) values to analyze the difference between the two methods. We observed that on spatial correlation metric (natural images: ours 78.1%, Shen et al. (2017) 76.1%) our method outperformed Shen et al. (2017) but on human judgment metric the results from Shen et al. (2017) were better compared to our method (natural images: ours 95.7%, Shen et al. (2017) 99.1%). In the method from Shen et al. (2017), they use a natural image prior that causes their reconstructions to look more natural and to outperform our method in terms of human judgment. We tried to introduce a natural -image prior through using a discriminator but the reconstructions did not appear as natural as compared to the results of Shen et al. (2017). However, in this work, our focus is not to propose a better reconstruction method, but to evaluate the potential of an end-to-end method to learn direct mapping from fMRI data to visual image.

### 3.2. Effect of dataset size

The results of the previous analyses show that it is possible to train the model with only 6,000 training samples from scratch. Therefore, we sought to investigate the effect of dataset size on the reconstruction quality. Furthermore, we attempted to check the possibility of improving the reconstruction quality by using more training samples.

We performed our analysis with an increasing training dataset from 120 to 6,000. We trained the model with 120, 300, 600, 1,500, 3,000, and 6,000 training samples and showed a qualitative comparison through the reconstructions (Figure 4A) and quantitatively through the pixel correlation and human judgment scores (Figure 4B). Through visual inspection of the reconstruction results in Figure 4A, we could observe that reconstruction quality improves on increasing the number of training samples. Pixel-wise correlation and human judgment scores (Figure 4B) also exhibit an increasing trend with increasing the number of training samples. The accuracy score improvement that follows with increasing number of training samples suggests that although we can obtain highly accurate reconstructions with only 6,000 training samples, there is still room for improvement, and better reconstruction quality could possibly be achieved given a larger dataset is available.

**Figure 4.**
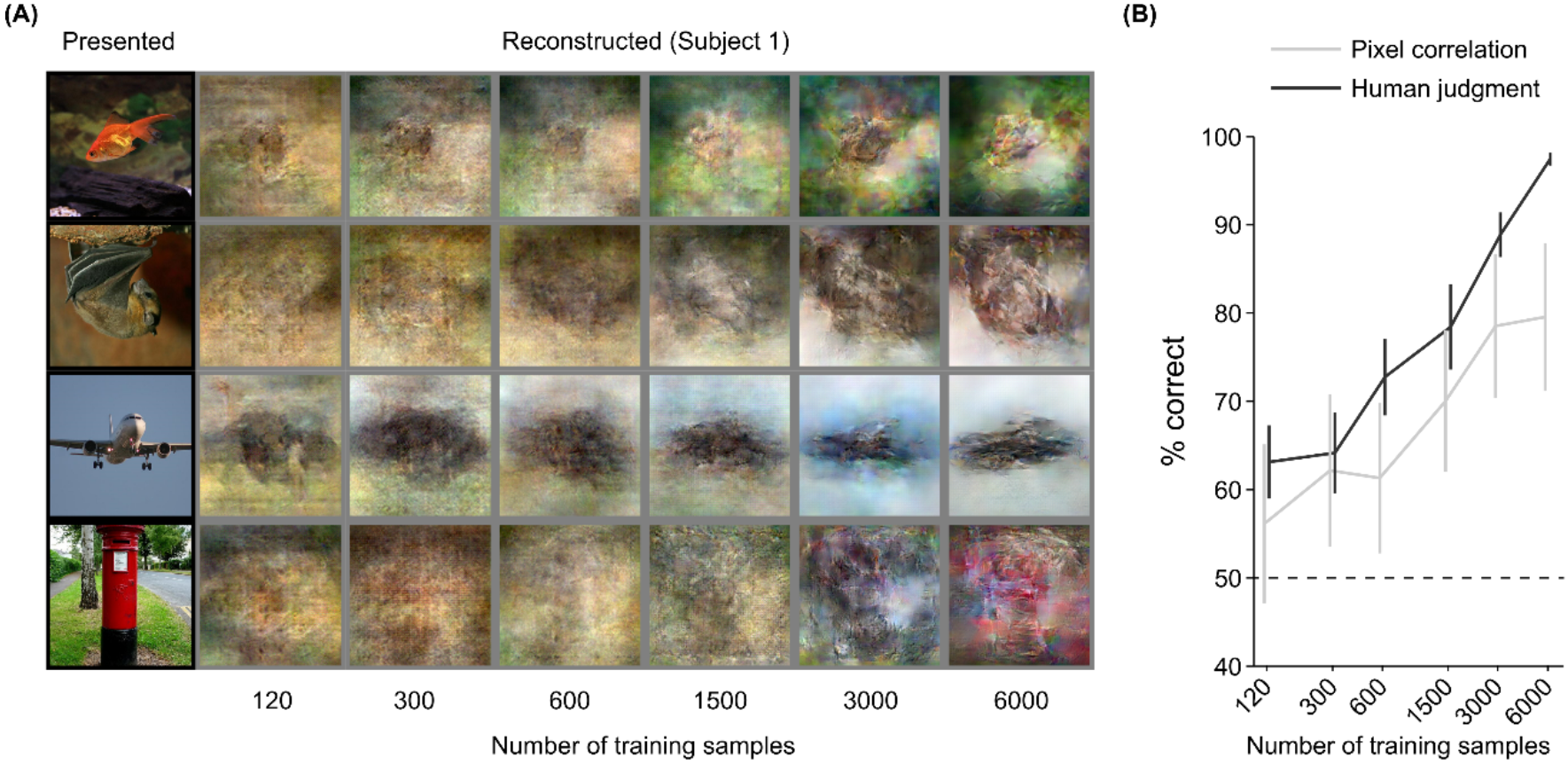
Effect of training dataset size. **(A)** Reconstruction from brain activity (Subject 1) using reconstruction models trained with different training dataset sizes. The presented images (in black frames) are shown in the first column. The corresponding reconstructed images (in gray frames) are shown to the right of each presented image (from left to right, the number of training samples increases). **(B)** Reconstruction accuracy in terms of percentage of correct pair-wise classification based on both pixel correlation and human judgment (error bars, 95% CI across samples; single subject (Subject 1), chance level, 50%).

### 3.3. Effect of loss functions: Ablation study

We performed an ablation study to understand the effect of different loss functions used in training the reconstruction model. We removed one loss function at a time and compared the reconstructions with those obtained using all three loss functions. Figure 5A shows the comparison results of the reconstructions obtained after removing one loss function at a time. On visually assessing the reconstruction results from Figure 5A, the reconstructions obtained from the model with all three loss terms show the best resemblance to the original stimulus images.

**Figure 5.**
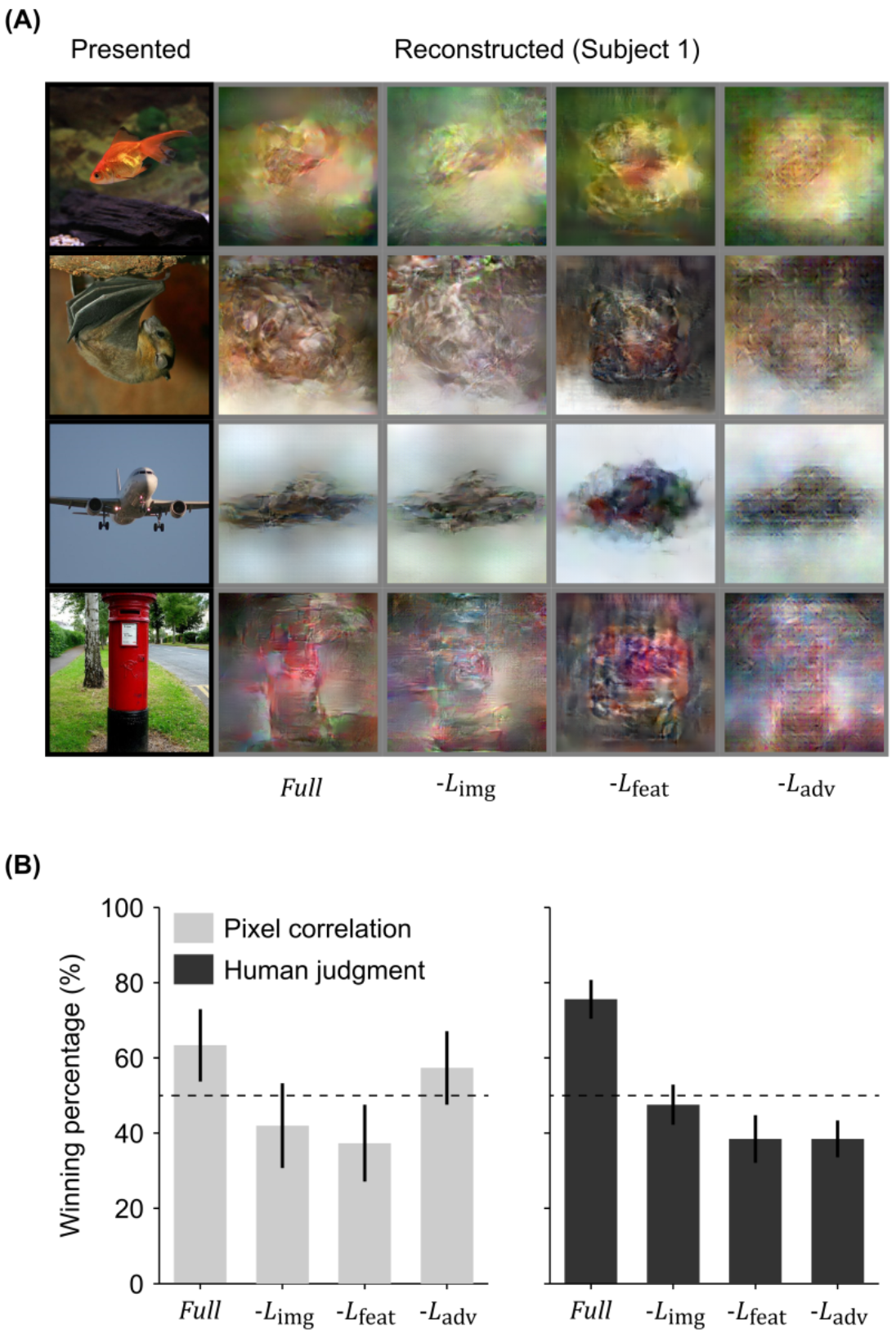
Ablation study of loss terms. **(A)** Reconstruction from brain activity (Subject 1) using the reconstruction model with some components of loss removed. The presented images (in black frames) are shown in the first column. The corresponding reconstructed images (in gray frames) obtained with different models are shown to the right of each presented image (from right to left, the model is: full reconstruction model (*Full*), with image loss removed (−*L*_img_), with feature loss removed (−*L*_feat_), and with adversarial loss removed (−*L*_adv_). **(B)** Reconstruction accuracy in terms of winning percentage of pair-wise classification based on both pixel correlation and human judgment (error bars, 95% CI across samples; single subject (Subject 1) chance level, 50%).

We observed that the difference in the human judgment and pixel correlation scores was not significant for the ablation study. So, to quantitatively compare the reconstruction quality for the ablation study we used the winning percentage as our criteria for comparison. The difference in winning percentage between the model optimized with all three loss terms and the model optimized with one loss term removed indicates the importance of the corresponding loss term. From Figure 5B, we can observe that the model trained with all three loss terms showed the highest winning percentage followed by the model where the loss in the image space is removed. The results demonstrate that the model trained with all three loss terms was preferred by the human raters as compared to the other models. Removing loss in the image space shows a similar drop for both of the winning percentage analyses (pixel correlation 21.3%, human judgment 28.0%) but the difference is not as pronounced as the other two loss functions. Removing feature loss shows the highest drop in the winning percentage for both spatial correlation (26.0%) and human judgment (37.2%). This demonstrates the importance of optimization in high dimensional feature space as it not only enhances the spatial details, but also makes the reconstruction more perceptually similar to its corresponding original stimulus image for human raters. Removing adversarial loss does not show much difference for spatial correlation criterion (6.0%), but, in the case of human judgment criterion, the difference is high (37.2%). This suggests that optimizing the adversarial loss forces the reconstruction to appear closer to natural image distribution.

## 4. Discussion

We have demonstrated that it is possible to learn a direct mapping function from fMRI activity in the visual cortex to the stimulus observed during perception. We showed this by performing an end-to-end training of a DNN model which reconstructs perceived stimuli from fMRI data. The reconstruction results on natural images obtained from the trained model show strong resemblance to the perceived stimuli in shape and in some cases color as well. Although trained only on natural images, the model generates accurate reconstructions of artificial shapes and alphabetical letters, thus showing good generalization comparable to Shen et al. (2017). We also demonstrated that the reconstruction quality improves as we increase the number of training samples and thus we believe that with more training samples we may be able to further improve the reconstruction accuracy.

We performed an ablation study by removing one loss function at a time to understand the importance of each loss term used for training the proposed model. The results showed that the model trained with all three loss terms achieved the best performance in terms of winning percentage. The removal of loss in image space resulted in similar change in the winning percentage calculated from behavioral experiments and spatial correlation scores. The removal of feature loss showed a significant drop in the winning percentage for both human ratings and spatial correlation, though the drop in human ratings was more pronounced. This implies that optimization in feature space enhances both spatial and perceptual similarity of the reconstructed image with the original stimulus. The removal of adversarial loss showed no significant drop in terms of spatial correlation but the drop in human rating result is quite high. This suggests that the addition of adversarial loss in the optimization process constrains the reconstructed image to be closer to training image distribution, leading to a significant difference in human ratings.

Earlier studies on decoding the stimulus in pixel space either search for a match in the exemplar set (Naselaris et al., 2009; Nishimoto et al., 2011) or try to obtain the exact reconstruction of the stimulus (Miyawaki et al., 2008; Wen et al. 2017; Güçlütürk et al. 2017; Shen et al. 2017; Han et al. 2017; Seeliger et al. 2017). In the exemplar matching methods, the visualization is limited to the samples in the exemplar set and hence these methods cannot be generalized to stimuli that are not included in the exemplar set. The reconstruction methods, however, are more robust to generalization to a new stimulus domain. Recent work from Shen et al. (2017) extended the reconstruction approach by capitalizing on multiple layers of DNN features, which were predicted from brain activity. They show that the decoders trained on only natural images can be successfully used to obtain reconstructions of artificial shapes and alphabetical letters.

DNN based reconstruction methods (Güçlütürk et al. 2017; Shen et al. 2017; Han et al. 2017, Seeliger et al. 2017) avoided directly training a DNN model for reconstruction. Instead, they used decoded features as an intermediate representation of the fMRI activity that was used as the input to a reconstruction module. This method is effective since the decoded features can easily be plugged into known image reconstruction/generation methods. It is also thought to be efficient given the lack of large-scale diverse fMRI datasets as compared to computer vision datasets used for end-to-end training of vision tasks. This makes it difficult to learn a direct mapping from brain activity to stimulus space without overfitting to the training dataset. Thus, learning this direct mapping from limited training samples was the main motivation of this work.

A potential advantage of the direct mapping is that it is free from constraints imposed by the pre-trained DNN and the features derived from a large scale image dataset. Even though the decoded features are correlated with the original image features, in Horikawa and Kamitani (2017) the maximum correlation coefficient on average was less than 0.5. So, we do not believe that information in the decoded features is the maximum visual information that can be extracted from the brain. Therefore, if enough training samples are available, a direct mapping may help in preventing this information loss.

In the present study, we demonstrated that it is possible to skip the intermediate step of feature decoding by an end-to-end approach, which allows us to learn a direct mapping from fMRI data to the perceived stimulus. Although the reconstructions obtained using the proposed method were highly accurate with good generalizability, our method could not outperform the method using decoded features on the available dataset. However, the results from the dataset size analysis suggest that the reconstruction quality could possibly be improved by increasing the size of the training dataset.

## Author contributions

Y.K. directed the study. G.S., K.D., and K.M. developed the reconstruction methods, G.S. and T.H. performed the experiments and analyses. K.D. and Y.K. wrote the paper.

## Acknowledgements

The authors thank Mohamed Abdelhack for valuable comments on the manuscript and Mitsuaki Tsukamoto for the help in computational environmental setting.

